# Susceptibility of wild canids to severe acute respiratory syndrome coronavirus 2 (SARS-CoV-2)

**DOI:** 10.1101/2022.01.27.478082

**Authors:** Stephanie M. Porter, Airn E. Hartwig, Helle Bielefeldt-Ohmann, Angela M. Bosco-Lauth, J. Jeffrey Root

**Author notes:** These senior authors contributed equally to this article (Angela M. Bosco-Lauth, J. Jeffrey Root).

## Abstract

Severe acute respiratory syndrome coronavirus 2 (SARS-CoV-2) has proven to be a promiscuous virus, capable of infecting a variety of different animal species, but much work remains in determining the susceptibility of common wildlife species to the virus. Here, we demonstrate that following experimental inoculation with SARS-CoV-2, red fox (*Vulpes vulpes*) become infected and can shed virus in oral and respiratory secretions. Conversely, experimentally challenged coyotes (*Canis latrans*) did not become infected or shed virus. Our results add red fox to the animal species known to be susceptible to SARS-CoV-2 and suggest that they may contribute to continued maintenance and transmission of the virus.

**Article Summary Line:** Experimental infection of red fox (*Vulpes vulpes*) and coyotes (*Canis latrans*) with SARS-CoV-2 revealed that red fox are susceptible to infection and can shed virus, while coyotes do not become infected.

## Introduction

Severe acute respiratory syndrome coronavirus 2 (SARS-CoV-2), the etiologic agent of the current coronavirus disease (COVID-19) pandemic, is believed to have been spilled over into the human population from a non-human mammalian source (*1*). SARS-CoV and MERS-CoV, two other betacoronaviruses that have caused outbreaks in humans in the early 21^st^ century, both likely originated in bats, with intermediate hosts believed to be masked palm civets (*Paguma larvata*) and dromedary camels (*Camelus dromedarius*), respectively (*2*). It is likely that SARS-CoV-2 also originated in bats, supported by the sequencing of a closely related betacoronavirus from horseshoe bats in China (*3*). Given that SARS-CoV-2 is likely a virus of zoonotic origin, and that over the course of the COVID-19 pandemic many animals have become infected with the virus, greater understanding of how animals could contribute to the epidemiology of SARS-CoV-2 will help direct further responses to controlling the pandemic.

Experimental infection studies have helped to expand our knowledge of the susceptibility of various animal species to SARS-CoV-2. Among members of the mammalian order Carnivora, experimental infections have revealed that domestic cats (*Felis catus*), domestic dogs (*Canis lupus familiaris*), ferrets (*Mustela putorius furo*), raccoon dogs (*Nyctereutes* procyonoides), and striped skunks (*Mephitis mephitis*) are susceptible to infection (*4-7*). Among those species, transmission of virus to naïve conspecifics has been experimentally demonstrated in cats, ferrets, and raccoon dogs (*4, 6, 8*). Domestic dogs appear to be minimally permissive to SARS-CoV-2, as experimental inoculations result in RT-PCR positive samples and low titer antibody responses, but no clinical disease or shedding of infectious virus (*4, 5*).

Natural infections with SARS-CoV-2, diagnosed by serology and/or PCR, have been reported in animals belonging to numerous carnivorous species. As of December 2021, cases in carnivores located in the United States include pets (domestic dogs, domestic cats, a ferret), captive animals in zoos [tigers (*Panthera tigris*), lions (*Panthera leo*), snow leopards (*Panthera uncia*), Asian small-clawed otters (*Aonyyx cinereus*), spotted hyenas (*Crocuta Crocuta*), a binturong (*Arctictis binturong*), a cougar (*Puma concolor*), a fishing cat (*Prionailurus viverrinus*), a Canada lynx (*Lynx canadensis*)], captive mink (*Neovision vision*) on farms, as well as free-ranging mink captured near farms (*9, 10*). Presumably, these animals contracted the virus from being in close proximity to humans (i.e., reverse zoonosis). In multiple instances, similar or completely identical viral sequences have been isolated from the owners and their pet dogs (*11-13*). In each of these instances, the epidemiologic timeline, in conjunction with sequence homology, suggests that human-to-animal transmission has occurred. Similar to experimentally infected dogs, the majority of pet dogs with confirmed SARS-CoV-2 infection are asymptomatic, but there have been multiple infected dogs with mild, non-specific clinical signs including coughing, sneezing, nasal discharge, dyspnea, diarrhea, and weakness (*12, 14-17*). A specialty veterinary hospital in the United Kingdom reported that concurrent to a wave of SARS-CoV-2 cases in humans, they observed an increase in the number of pet dogs and cats presenting with signs of myocarditis (*18*). Opportunistic sampling of pets with cardiac disease at this hospital revealed that 6 of 11 animals tested via RT-PCR or serology had been infected with SARS-CoV-2 (*18*).

The ability of SARS-CoV-2 to infect domestic dogs, in addition to several other species of carnivores, suggests that other members of the canid family may be susceptible to infection. Wild canids, such as red fox (*Vulpes vulpes*) and coyotes (*Canis latrans*), are of particular interest given how widespread these animals are distributed, their often close proximity to humans, and the fact that they may prey and scavenge upon, or otherwise interact with species demonstrated to be susceptible to SARS-CoV-2, including felids, skunks, rodents, and white-tailed deer (*19, 20*). Both red fox and coyote populations are widely distributed and can be found in environments ranging from rural to urban. These species have become particularly adapted to urban environments; red fox can be found in major cities worldwide, and coyotes have been seen in every large city in the continental United States (*21, 22*). Studies of urban adapted coyotes have found that they are less likely to flee from humans, more likely to engage in conflicts with pets, and more likely to prey on pet dogs and cats than coyotes in rural environments, all behaviors that could increase the coyotes’ risk of being exposed to SARS-CoV-2 (*22, 23*).

Fox have been included in modeling efforts and serosurveillance studies aiming to predict animal hosts of SARS-CoV-2. Structural analysis of the ACE2 receptor in various animal species predicted red fox ACE2 to have the ability to bind SARS-CoV-2, with different models predicting low, medium, or high affinity binding (24-26). Thus far, surveillance studies to ascertain the prevalence of SARS-CoV-2 in wildlife have infrequently screened wild canids. A serosurvey conducted in China in early 2020 failed to show any seropositive fox (species not specified) in a cohort of 89 wild animals (*27*). A serosurvey conducted in Croatia from June 2020 to February 2021 revealed 6 of 204 (2.9%) of tested red fox to be ELISA positive for SARS-CoV-2 antibodies (*28*). However, despite being repeatably positive with ELISA, these results could not be confirmed with a surrogate virus neutralization test (sVNT). These two surveys were both conducted relatively early during the COVID-19 pandemic, and do not exclude the possibility that red fox may become infected with and develop an immune response to SARS-CoV-2, only that spillover had not occurred in the examined animals in those particular locations at the time of the studies. As numerous carnivore species have proved to be susceptible to SARS-CoV-2, ascertaining the susceptibility of other wild carnivores to the virus, especially those species that are closely associated with humans, is a crucial step in understanding the role that wildlife may play in maintaining and transmitting SARS-CoV-2. The objective of this study was to assess two species of wild canids – red fox and coyotes – for susceptibility to infection with SARS-CoV-2.

## Method

### Animals

Captive reared, juvenile (3-5-month-old), mixed sex red fox (3 female, 3 male) and coyotes (3 female, 1 male) were evaluated for susceptibility to SARS-CoV-2. Animals were individually housed in an animal biosafety level-3 (ABSL-3) facility at Colorado State University (CSU). Animal work was approved by CSU and National Wildlife Research Center Institutional Animal Care and Use Committees. All animals had access to food and water *ad libitum*. Red fox were maintained on Fromm dog food, while coyotes were maintained on Mazuri exotic canine diet; both species were supplemented with fresh fruit, mealworms, and eggs. Temperature-sensing microchips (Bio-thermo Lifechips, Destron Fearing) were subcutaneously implanted in all animals.

### Virus

SARS-COV-2 strain WA1/2020WY96 obtained from BEI Resources was passaged twice in Vero E6 cells, and stock virus harvested in Dulbecco’s Modified Eagle Medium (DMEM) with 5% fetal bovine serum and antibiotics was frozen at -80°C. Double overlay plaque assay on Vero cells, with plaques counted 72 hours later, was then performed to titrate virus stock and determine plaque forming units (pfu) per mL of stock (*4*).

### Virus challenge

Animals were lightly anesthetized with 6-11 mg/kg ketamine hydrochloride (Zetamine) and 0.6-1.1 mg/kg xylazine given intramuscularly. A baseline body temperature, weight, oral swab, and blood sample was collected from each animal prior to inoculation. Virus diluted in phosphate buffered saline (PBS) was instilled into the nares of each animal (500 µl per nare, for a total volume of 1 mL). Animals then were observed until fully recovered from anesthesia. Virus back-titration on Vero cells was then immediately performed; each animal was confirmed to have received between 5.1-6.0 log_10_ pfu of SARS-CoV-2.

### Sampling

Animals were lightly anesthetized using the aforementioned protocol for all sample collections. Oropharyngeal swabs were collected on days 1, 2, 3, 5, 7, and 14 days post-inoculation (dpi). Swabs were placed in 1 mL tris-buffered MEM with 1% bovine serum albumin, supplemented with gentamycin, amphotericin B, polymyxin B, and penicillin/streptomycin (BA-1). Nasal flushes were performed on days 1, 2, 3, 5, and 7 dpi, by instilling 1 mL of BA-1 into the nares and collecting the nasal discharge that was sneezed or dripped out onto a sterile petri dish. One-half of the animals (3 red fox, 2 coyotes) were euthanized and necropsied at 3 dpi to evaluate tissues for acute viral burden and pathological changes. The following tissues were routinely collected at necropsy for virus isolation and formalin fixation: nasal turbinates, trachea, lung, heart, liver, kidney, spleen, and small intestine. The remaining animals were maintained until 28 (red fox) or 30 (coyote) dpi in order to screen for serologic responses. These animals had blood collected at 7 (red fox) or 8 (coyote), 14, 21, and 28 (red fox) or 30 (coyote) dpi. Animals were then euthanized, necropsies performed, and the previously listed tissues were collected and fixed in formalin.

### Clinical Observations

All animals were assessed daily for attitude and signs of clinical disease, including lethargy, anorexia, ocular discharge, nasal discharge, sneezing, coughing, and dyspnea. To assess for weight loss and pyrexia, the weights and temperatures of all animals were measured each time an animal was handled.

### Virus Isolation

Viral isolation was performed on all oral swab, nasal flush, and 3 dpi tissue samples by double overlay plaque assay on Vero cells (*4*). Serial ten-fold dilutions of samples in BA-1 were prepared, and 100 µL was used to inoculate confluent monolayers of Vero cells grown in 12 well plates. Plates were then incubated for one hour at 37°C, and the cells overlaid with 0.5% agarose in MEM with 2% fetal bovine serum, antibiotics, and antifungal agents. A secondary overlay with neutral red dye was added at 24 hours, and plaques were counted at 48-72 hours. Viral titers were reported as log_10_ plaque-forming units (pfu) per mL for oral swab and nasal flush samples, and per gram for tissues.

### Serologic Analysis

Plaque reduction neutralization assays were performed on all serum samples (*4*). Serum samples were heat inactivated for 30 minutes at 56°C. Serial two-fold dilutions in BA-1 were prepared (starting at 1:20), with the exception of pre-inoculum serum, which were prepared at 1:10. These dilutions were aliquoted onto 96 well plates, and an equal volume of virus was added to each well. Plates were then incubated at 37°C for one hour. Following the incubation, the samples were plated onto confluent monolayers of Vero cells and a double overlay performed, as described above for plaque assays. Antibody titers were recorded as the reciprocal of the highest dilution in which >80% of virus was neutralized.

### qRT-PCR

Plaques were picked from culture plates of all positive animals to confirm the presence of SARS-CoV-2 RNA. RNA was extracted from plaque picks, as well as oral swabs from plaque assay negative animals, using QiaAmp Viral RNA Mini Kits (QIAGEN). RT-PCR was then performed using the E-sarbeco primer probe sequence described elsewhere and the Qiagen QuantiTect Virus Kit (*29*). The thermal cycling protocol consisted of 20 minutes at 50°C for reverse transcription, 95°C for 5 minutes, followed by 50 cycles of 95°C for 15 seconds and 58°C for 45 seconds.

### Histopathology

Animal tissues were fixed in 10% neutral buffered formalin for 14 days, then processed for paraffin embedding and sectioning. Slides were stained with hematoxylin and eosin, and read by a veterinary pathologist.

## Results

### Clinical Observations

Neither weight loss nor elevated temperatures were observed in any animals during the course of the study. On 4 dpi, one red fox was observed to be lethargic, and on 6 dpi all three remaining red fox were lethargic and sneezing. No other behavioral changes or clinical signs of disease were seen in any of the animals at any other time point.

### Virus Isolation

All six of the red fox shed infectious virus both orally (Figure 1) and nasally (Figure 2) starting at 1 dpi. A majority of the red fox were still shedding virus at 3 dpi (4/6 oral, 5/6 nasal), with all shedding resolved by 5 dpi. Of the tissues from the fox euthanized at 3 dpi, infectious virus was isolated from the nasal turbinates of two of three animals, and not from any other tissues. Infectious virus was not isolated from any of the oral swabs, nasal flushes, or tissues collected from any of the coyotes.

**Figure 1.**
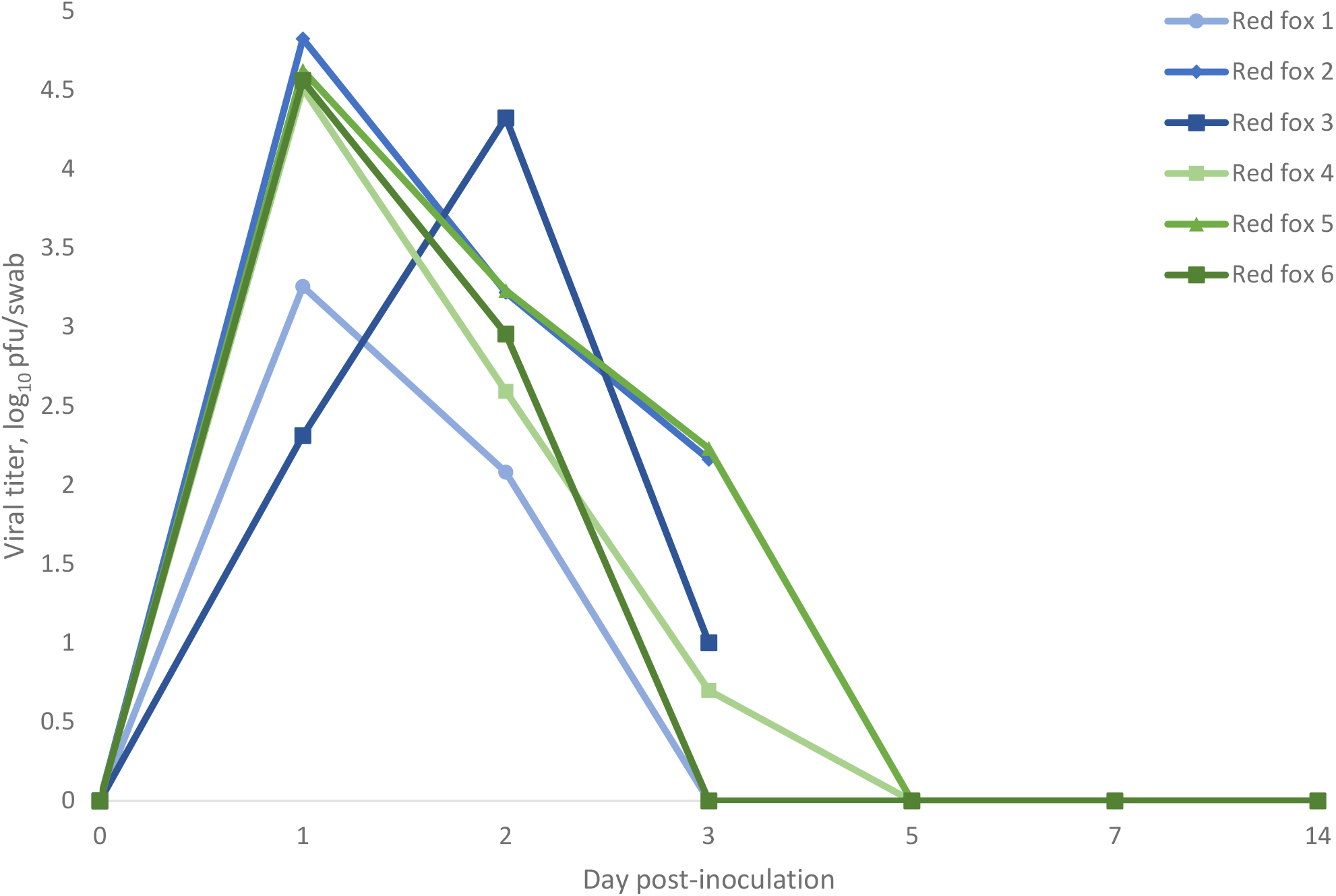
Oropharyngeal shedding of SARS-CoV-2 by red fox as detected by plaque assay. Values are expressed as log_10_ pfu/swab. Red fox 1, 2, and 3 were euthanized at 3 dpi.

**Figure 2.**
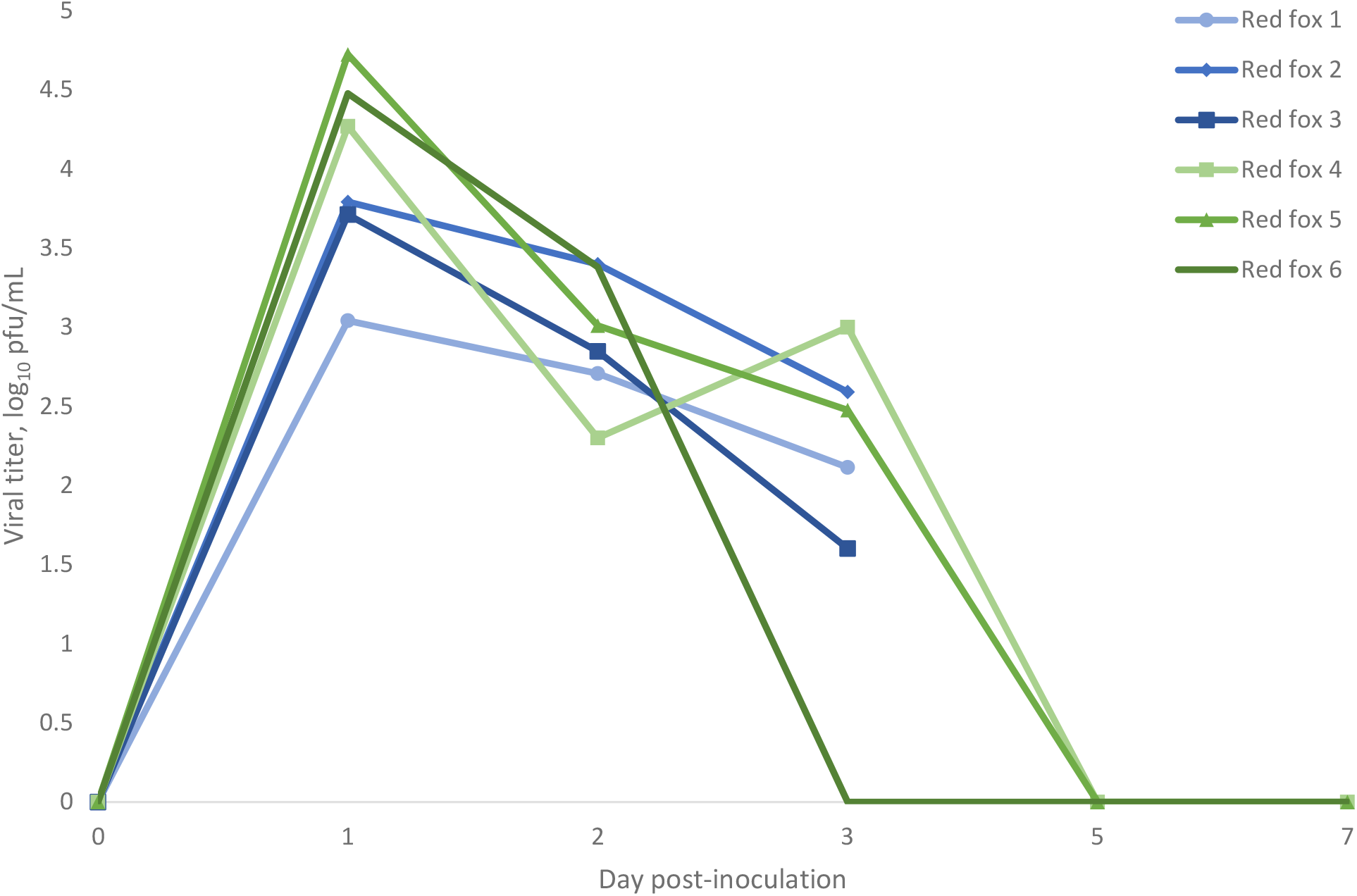
Nasal shedding of SARS-CoV-2 by red fox as detected by plaque assay. Values are expressed as log_10_ pfu/mL. Red fox 1, 2, and 3 were euthanized at 3 dpi.

### Serology

Prior to inoculation, all animals were seronegative against SARS-CoV-2. None of the animals euthanized at 3 dpi developed an acute neutralizing antibody response. All of the red fox held until 28 dpi showed a neutralizing antibody response beginning at 7 dpi, with peak titers (1:80 or higher) reached at 14 dpi (Table 1). None of the coyotes seroconverted.

**Table 1.** Antibody titers (PRNT80) for red fox experimentally infected with SARS-CoV-2

| **Animal** | **Pre-infection** | **7 dpi\*** | **14 dpi** | **21 dpi** | **28 dpi** |
| --- | --- | --- | --- | --- | --- |
| **Red fox 4** | 0 | 20 | 160 | 80 | 80 |
| **Red fox 5** | 0 | 20 | 80 | 80 | 80 |
| **Red fox 6** | 0 | 20 | 320 | 160 | 320 |

Table footnotes: \*dpi = days post-inoculation

### Pathology

On necropsy, no gross lesions were observed. None of the fox tissues evaluated had histopathologic lesions attributable to SARS-CoV-2. Tissues from the coyotes were not assessed as they neither shed infectious virus nor seroconverted.

### qRT-PCR

All red fox were confirmed via RT-PCR to have shed SARS-CoV-2. The 2 dpi oral swabs from all four coyotes were positive for SARS-CoV-2 RNA, albeit with high CT values (range 32-35). All other coyote oral swabs were negative for SARS-CoV-2.

## Discussion

The COVID-19 pandemic continues to present a challenge globally. An understanding of how non-human animals may contribute to the epidemiology of SARS-CoV-2 is key in the continued response to curb the pandemic, as well as understanding the impacts of the virus on animal health. Both experimental and natural infections have contributed to our knowledge of the wide range of mammalian species that can be infected with SARS-CoV-2. In the vast majority of cases, non-human animals seem to experience subclinical or mild disease following infection with SARS-CoV-2. There are exceptions – infected mink, large felids in zoos, and the occasional pet dog or cat have presented with signs of a respiratory infection, including coughing, sneezing, ocular/nasal discharge, and increased respiratory effort (*30-32*). More severe disease has also been observed, with a report out of the United Kingdom hypothesizing that SARS-CoV-2 infection may be linked to myocarditis in pet dogs and cats (*18*), and mink farms reporting increased mortality during SARS-CoV-2 outbreaks (*30*). Thus far, only a few animal species have demonstrated the ability to transmit virus to naïve conspecifics (deer mice, cats, ferrets, raccoon dogs, and white-tailed deer), and mink are the only animals that have definitively infected humans (*33-35*). In this study, we add to the present knowledge base of SARS-CoV-2 in animals by describing mild clinical disease and viral shedding in an additional species, the red fox, while describing lack of susceptibility in coyotes.

At present, the COVID-19 pandemic has been driven by human-to-human transmission of SARS-CoV-2, but animal species that are susceptible to infection with the virus represent a niche for viral maintenance, and could potentially serve as a source for viral spillback into the human population, as has already been the case on mink farms. Therefore, peridomestic species (animals occupying habitats in close proximity to humans) are of particular interest as they presumably run the greatest risk of contracting the virus from people.

We demonstrated that red fox are highly susceptible to infection with SARS-CoV-2. All red fox in this study shed infectious virus both orally and nasally for at least three days. Each of the red fox held for 28 days displayed mild, self-resolving, clinical signs including lethargy and sneezing, and developed neutralizing antibody responses beginning 7 dpi, and persisted for the duration of the study. These antibody titers are similar to what is seen in experimentally infected domestic dogs (*4*). Despite this evidence of mild infection, no histopathological changes attributable to SARS-CoV-2 were observed in red fox tissues; the lack of lesions in the heart is notable given reports of myocarditis in pet dogs infected with SARS-CoV-2 (*18*). Conversely, coyotes appear to not be susceptible to infection with SARS-CoV-2, as none of the animals in the study shed detectable virus or seroconverted following challenge. While coyote oral swabs were positive for viral RNA on 2 dpi, this was not associated with isolation of infectious virus, and likely represents either residual inoculum or an infection below the limit of detection. Hence, coyotes are unlikely to be competent hosts for SARS-CoV-2.

While small sample sizes and direct intranasal inoculation with a high dose of virus are limitations of experimental challenge studies like this one, these studies do provide important information that can inform continued surveillance and control efforts. The animals found to be susceptible to natural infection with SARS-CoV-2 have reflected results from experimental challenge studies, so it is reasonable to assume that our results can be extrapolated. Therefore, special attention should be paid to red fox when considering wildlife species that may serve as reservoir hosts for SARS-CoV-2. We demonstrated that SARS-CoV-2 infected red fox shed infectious virus for multiple days in both oral and nasal secretions, which suggests that red fox could contribute to onward transmission of the virus. As red fox commonly consume, via hunting or scavenging, other species that are susceptible to infection with SARS-CoV-2, including rodents and white-tailed deer (*20*), predator-prey interactions and scavenging may serve as avenues for inter-species transmission.

Should wildlife species such as red fox become established maintenance hosts of SARS-CoV-2, consequences could include impacts on animal health, development of novel viral variants, and spillback into the human population. Caution should be taken when interacting with susceptible wildlife species in order to prevent transmission events. To best curtail the COVID-19 pandemic, we should understand all that may contribute to the continued transmission of SARS-CoV-2, and potential maintenance and transmission of the virus by wildlife species could be an important component of SARS-CoV-2 epidemiology.

## Acknowledgements

This work was supported by internal funding from Colorado State University and the U.S. Department of Agriculture, Animal and Plant Health Inspection Service. We are very grateful to Julie Young (USDA) and Stacey Brummer (USDA) for providing training and the captive-bred coyotes used in this study, and to USDA-APHIS-WS-NWRC animal care staff for assistance with animal care. Special thanks to Jeremy Ellis (USDA) for invaluable technical assistance, and to Richard Bowen (Colorado State University) for guidance, consulting, and facilities support.

## Biographical sketch

Stephanie Porter is an APHIS Science Fellow with the National Wildlife Research Center at the United States Department of Agriculture. Her research interests include the pathogenesis and transmission of infectious disease.

